## Supplementary figures and images for "CryoEM structures of the human CLC-2 voltage gated chloride channel reveal a ball and chain gating mechanism"

### Figure 6 source data 2

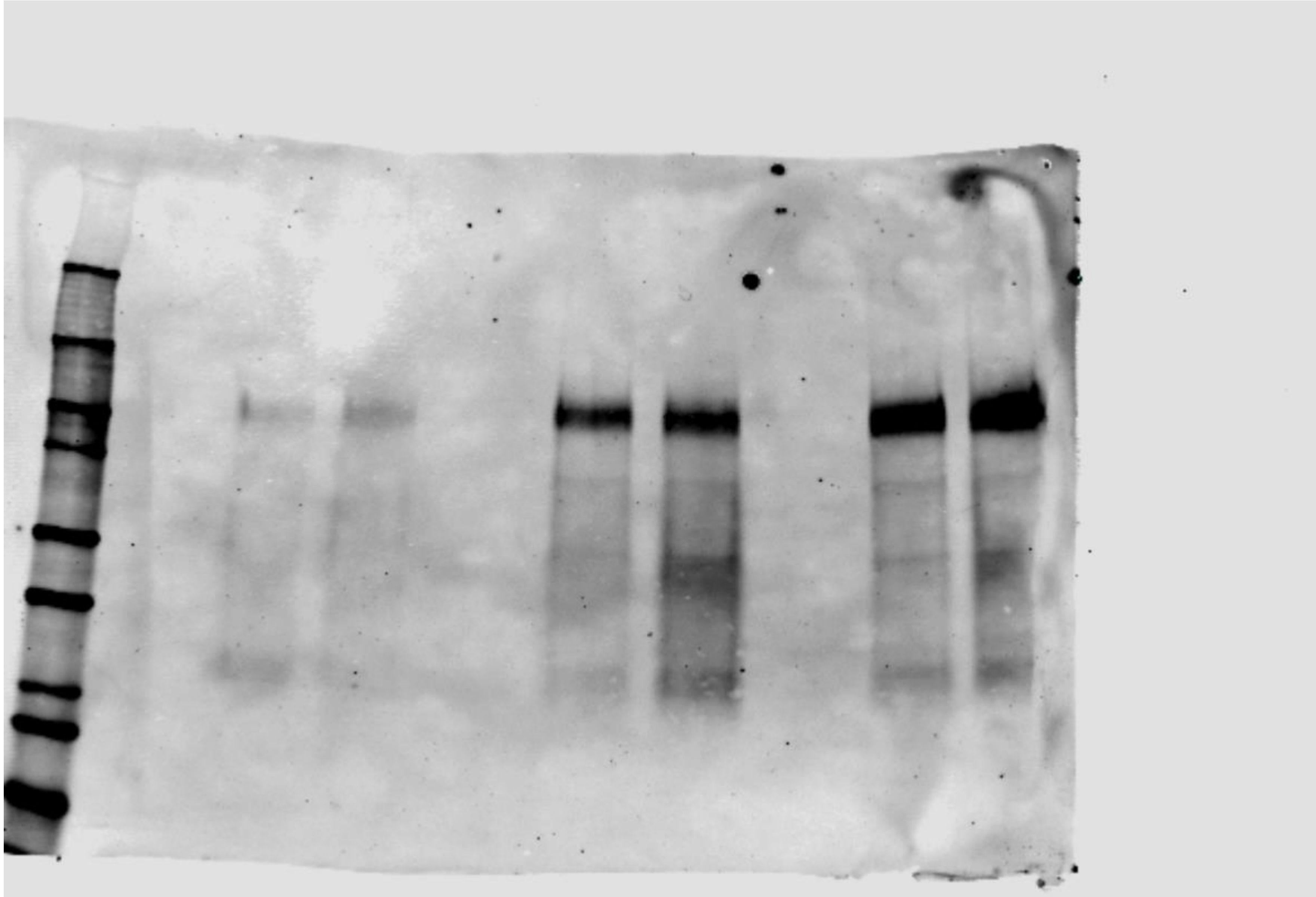
